# Functional and diffusion MRI reveal the functional and structural basis of infants’ noxious-evoked brain activity

**DOI:** 10.1101/2020.04.28.065730

**Authors:** Luke Baxter, Fiona Moultrie, Sean Fitzgibbon, Marianne Aspbury, Roshni Mansfield, Matteo Bastiani, Richard Rogers, Saad Jbabdi, Eugene Duff, Rebeccah Slater

## Abstract

Understanding the neurophysiology underlying pain perception in infants is central to improving early life pain management. In this multimodal MRI study, we use resting-state functional and white matter diffusion MRI to investigate individual variability in infants’ noxious-evoked brain activity. In an 18-infant nociception-paradigm dataset, we show it is possible to predict infants’ cerebral haemodynamic responses to experimental noxious stimulation using their resting-state activity across nine networks from a separate stimulus-free scan. In an independent 215-infant Developing Human Connectome Project dataset, we use this resting-state-based prediction model to generate noxious responses. We identify a significant correlation between these predicted noxious responses and infants’ white matter mean diffusivity, and this relationship is subsequently confirmed within our nociception-paradigm dataset. These findings reveal that a newborn infant’s pain-related brain activity is tightly coupled to both their spontaneous resting-state activity and underlying white matter microstructure. This work provides proof-of-concept that knowledge of an infant’s functional and structural brain architecture could be used to predict pain responses, informing infant pain management strategies and facilitating evidence-based personalisation of care.

## Introduction

Newborn infants routinely undergo numerous painful procedures as part of standard clinical care shortly after birth during their stay in hospital ^1^. Their lack of verbal communication, brief extra-uterine medical history, and ambiguity in the behavioural and physiological responses that underpin infant pain scales ^2^, lead to a high degree of uncertainty in clinical decision-making related to the treatment of infant pain. Understanding and anticipating an individual infant’s response to nociceptive input would advance efforts of personalised pre-emptive pain minimisation in this vulnerable population. In the experimental setting, a multitude of complementary behavioural, physiological, and neural measures are used in an attempt to quantify infant pain and pain sensitivity, with a high degree of individual variability observed across all modalities ^3–6^. In this study, we focus on newborn infants’ cerebral haemodynamic responses to experimental nociceptive input recorded using functional magnetic resonance imaging (fMRI). We test whether infants’ response amplitudes can be predicted from their resting-state brain activity and whether the amplitudes are associated with underlying white matter microstructure. The inherent limitation of small sample sizes in infant fMRI pain studies is mitigated by identifying consistent findings in a large independent age-matched sample from the Developing Human Connectome Project (dHCP) dataset (http://www.developingconnectome.org).

A high degree of correspondence between resting-state and task-related brain activities has been observed in adult fMRI studies ^7,8^. In adults, fMRI-recorded resting-state brain activity has been observed to be a distinguishing feature of an individual’s brain functionality ^9^, and has been used to predict individuals’ task-related brain activity under both experimental ^10^ and clinical conditions ^11^. While analogous studies have not been conducted in the newborn infant population, large-scale resting-state networks are detectable using fMRI from birth and appear to correspond to adult canonical resting-state and task-response networks ^12,13^, suggesting a similar correspondence could exist at this early stage of development. Using a cohort of 18 healthy newborn infants, we replicate large-scale resting-state networks that have previously been characterised in an independent age-matched subset of the dHCP dataset ^14^. The amplitudes of spontaneous activity of these networks were then used to predict the infants’ cerebral haemodynamic response amplitudes in response to an experimental nociceptive stimulus, in a cross-validated manner.

Previous studies in infants have demonstrated the sensitivity of infant noxious-evoked cerebral activity to sleep state ^15^ and physiological stress ^16^. To disambiguate temporally stable trait effects, arguably of higher relevance for clinical pre-emptive decision making, from temporally transient state effects, we assessed the correlation between infants’ nociceptive haemodynamic response amplitude and underlying white matter microstructure using diffusion MRI (dMRI) data. These white matter microstructural features will reflect the integrity of developing structural connectivity, which constrains infants’ noxious-evoked responses. Due to the dHCP dataset’s larger numbers, we used it to explored possible structure-function relationships across multiple white matter tracts for three microstructural parameters: mean diffusivity (MD), fractional anisotropy (FA), and mean kurtosis (MK). Predicted noxious-response amplitudes were generated from the dHCP infants’ resting-state data using the resting-state-based prediction model trained in our 18-infant nociception-paradigm dataset. Structure-function associations identified in the dHCP dataset were subsequently tested and validated in our nociception-paradigm dataset. Within the dHCP dataset, we found robust statistically significant negative correlations between (predicted) noxious-response amplitudes and the white matter MD of five bilateral tracts, suggesting infants with larger responses had more structurally mature connectivity. This negative correlation between (observed) noxious-response amplitudes and MD was directly confirmed within the nociception-paradigm dataset. This structure-function relationship, consistently identified in two independent datasets, suggests the infants’ haemodynamic responses are dependent on specific white matter microstructural features, likely white matter myelination or fibre packing density, and thus are temporally stable trait effects.

This work provides new insight into the neurophysiological basis for normative variability in the cerebral response to nociceptive input in a group of healthy newborn infants. A nociception-related neural structure-function relationship is revealed, and tight coupling between an infant’s resting-state and noxious-response neural activities provides proof-of-concept that an infant’s resting-state brain activity during periods which are free of nociceptive input can be used to make accurate predictions about their brain response to nociceptive stimuli.

## Results

### Infants displayed wide variability in haemodynamic response amplitude to nociceptive input

We quantified the change in brain activity evoked by a mild experimental noxious stimulus to the foot in 18 healthy newborn infants (Figure 1). The cerebral haemodynamic response to the 128 mN pinprick was highly variable between infants, and included both negative (3 of 18 infants) and positive (15 of 18 infants) blood oxygen level dependent (BOLD) responses (Figure 1 heat maps). Summarising each infant’s noxious-response map relative to the group average response map, the relative response amplitudes ranged from -0.87 to 5.60 (Figure 1 scalar values).

**Figure 1:**
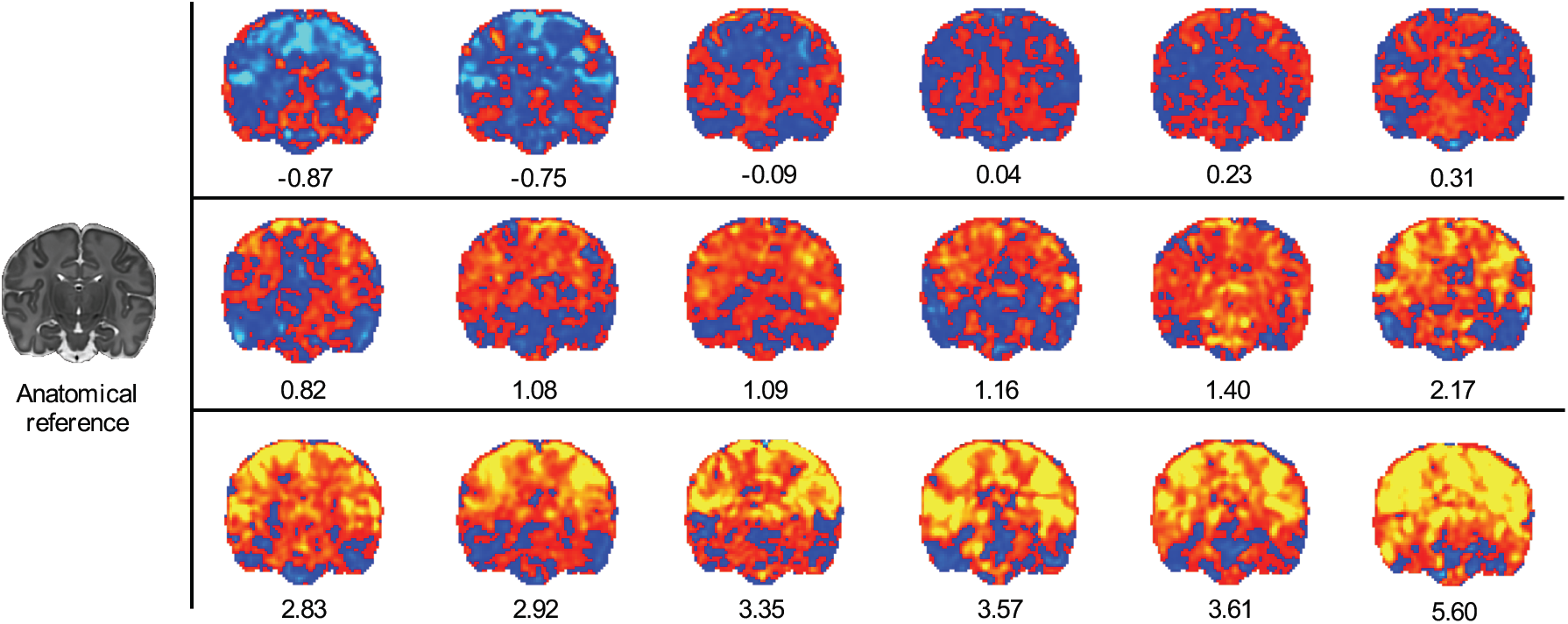
Newborn infants’ noxious-response amplitudes. A noxious-response BOLD activity map is presented for each infant (n=18) and ordered according to the response amplitude relative to the group average. The maps are general linear model regression parameter maps i.e. effect size maps. The scalar value presented below each map is a summary measure that represents the whole-brain noxious-response amplitude relative to the group average. It is calculated by spatially regressing the group-average noxious-response map onto each individual infant’s noxious-response map. Red-yellow indicates positive values and blue-cyan indicates negative values. The anatomical reference (left) provides structural detail for orientation. All noxious-response maps are displayed at this slice position.

In each infant, the noxious-evoked BOLD response was well fit by the infant double gamma haemodynamic response function (HRF) for both non-negligible positive and negative response amplitudes (Supplementary Information: *Noxious-response HRF fit assessment*). There were no obvious signs of gross artefactual errors, such as head motion-related spikes or variable response latencies, suggesting that the HRF-estimated noxious-response amplitudes reflect physiologically meaningful differences in the cerebral haemodynamic response amplitude to the noxious input.

### Nine resting-state networks were replicable across the nociception-paradigm and dHCP datasets

In the same cohort of 18 infants, nine resting-state networks were robustly identified from separate resting-state scans using probabilistic functional mode analysis ^17,18^ (Figure 2). These included three sensory and motor networks (two visual, two auditory, and two somatomotor networks) and three cognitive networks (default mode, dorsal attention, and executive control networks). To consider a network robust and suitable for inclusion in the subsequent analysis, networks needed to be consistent across both the nociception-paradigm cohort of 18 infants (Figure 2 top row) and a large independent cohort of 242 age-matched infants that were collected as part of the dHCP and analysed using the same analytical approach (Figure 2 bottom row). Matched networks were highly consistent between datasets with spatial Pearson correlation coefficients between unthresholded maps ranging from 0.63 to 0.90 (mean = 0.78) (Figure 2 scalar values).

**Figure 2:**
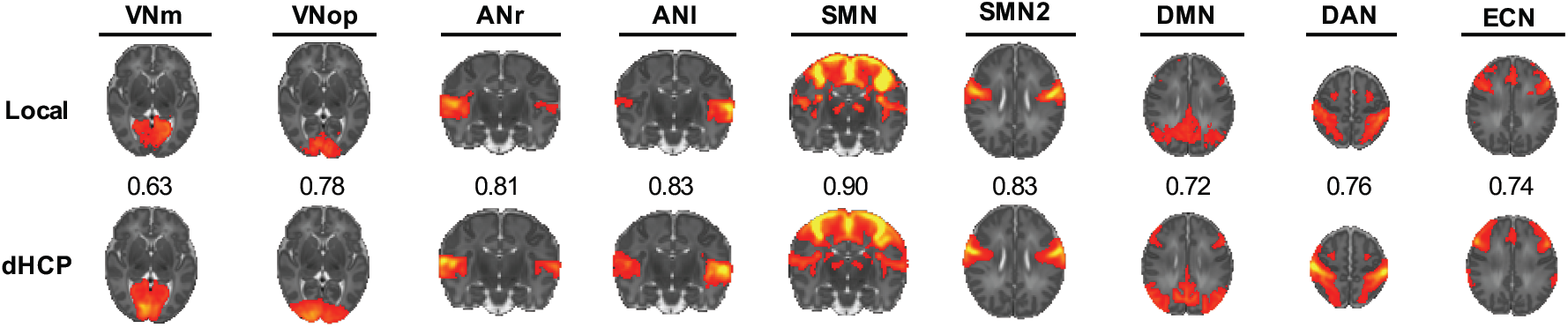
Nine resting-state networks replicated across two independent datasets. Each resting-state network map is a thresholded group-level probabilistic functional mode (PFM) map identified in the locally collected 18-infant nociception-paradigm dataset (top row, Local) and the 242-infant dHCP dataset (bottom row, dHCP). These PFM posterior probability maps are thresholded to highlight qualitative correspondence. The scalar value shown between matched maps is the spatial Pearson correlation coefficient between unthresholded maps highlighting quantitative correspondence. Abbreviations: VNm = medial visual network; Vnop = occipital pole visual network; ANr = right auditory network; ANl = left auditory network; SMN = somatomotor network; DMN = default mode network; DAN = dorsal attention network; ECN = executive control network.

### Resting-state network amplitudes predicted noxious-response amplitudes

Infants’ whole-brain noxious-response amplitudes (Figure 1 scalar values) were predicted from their task-free resting-state network amplitudes with statistically significant prediction accuracy (Figure 3 and Table 1 Resting state). The resting-state network amplitudes were quantified using multiple regression of the nine dHCP networks (Figure 2) onto each infant’s resting-state data, and each resulting network timeseries summarised as an amplitude using the median absolute deviation, which ensures robustness to outliers. Using a support vector regression (SVR) model, predictions were generated in a leave-one-out cross-validation (LOO-CV) approach, including cross-validated confound regression for several confound variables. Measures of prediction accuracy (Table 1) were tested for statistical significance using permutation testing.

**Table 1:** Noxious-response amplitude prediction accuracies. Each row contains results for a specific set of predictors. Each column contains prediction accuracy assessed using a specific metric: RMSE = root mean squared error; R^2^ = coefficient of determination (sums-of-squares formulation); R_Sp_ = Spearman’s rank correlation coefficient. P-values are presented in parentheses. * = statistically significant.

| | RMSE | $R^2$ | $R_{sp}$ |
| --- | --- | --- | --- |
| Resting state | 1.55* (0.004) | 0.64* (0.004) | 0.77* (0.003) |
| Confounds | 2.46 (0.60) | 0.081 (0.60) | 0.14 (0.40) |
| Clinical variables | 2.71 (0.47) | -0.12 (0.47) | 0.0031 (0.51) |

**Figure 3:**
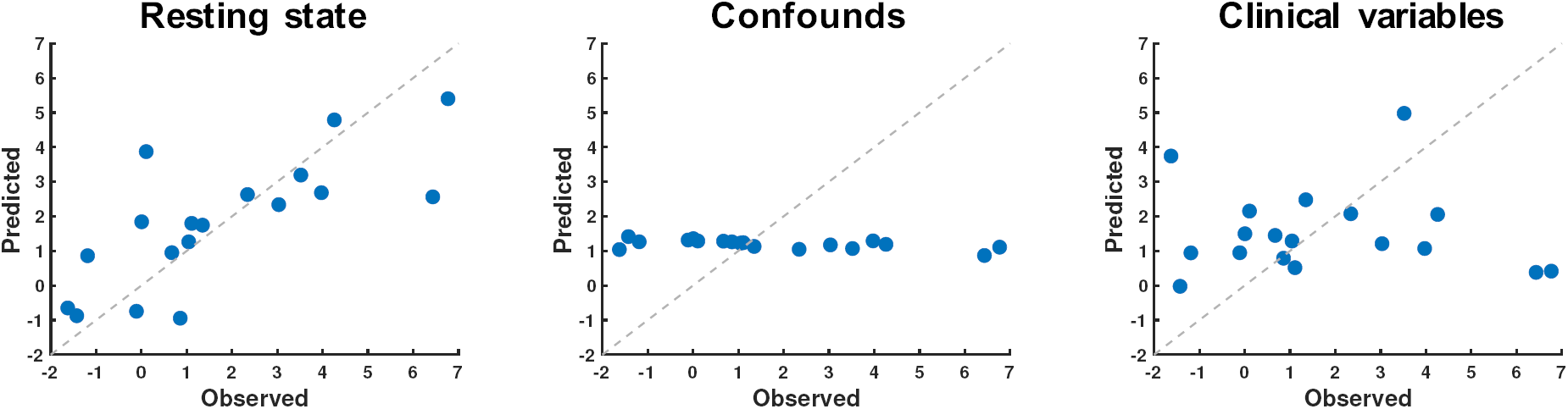
Predicting noxious-response amplitudes from non-noxious data. For all plots, each blue dot represents an out-of-sample cross-validated prediction for a single infant (n=18), and the dashed grey line is the y=x line along which perfect predictions would lie. The x-axis is the observed noxious-response amplitude (after cross-validated confound regression), and the y-axis is the predicted noxious-response amplitude. Predictions were generated based on three sets of predictors: (left) the resting-state network amplitudes; (middle) resting-state imaging confounds, which included head motion, CSF amplitude, and white matter amplitude; and (right) clinical variables, which included age (GA, PMA, and PNA), birth weight, TBV, and sex.

Three resting-state imaging confounds, which included head motion and cerebrospinal fluid (CSF) and white matter amplitudes, were additionally tested but were not predictive of the infants’ noxious-response amplitudes (Figure 3 and Table 1 Confounds). Similarly, six non-fMRI variables (henceforth named clinical variables), which included postmenstrual age (PMA), gestational age (GA), postnatal age (PNA), birth weight, total brain volume (TBV), and sex, were tested and were not predictive (Figure 3 and Table 1 Clinical variables). For both the resting-state imaging confounds and the clinical variables, predictions were centred on the LOO-CV training set mean noxious-response amplitudes (Figure 3 Confounds and Clinical variables). The lack of association between the noxious-response amplitudes and the resting-state imaging confounds suggested that the predictive capacity of the resting-state network amplitudes was not mediated by undesirable features of resting-state data, but rather appeared to be mediated by the correspondence between an individual infant’s resting-state and noxious-evoked brain activities. These brain function similarities could not be explained by the infant’s age, birth weight, brain volume, or sex.

Additionally, the number of resting-state network timeseries outliers (an indicator of resting-state network timeseries quality) was assessed and found to be unrelated to infants’ noxious-response amplitudes (Supplementary Information: *Resting-state network timeseries outlier assessment*). Finally, univariate correlation analyses between noxious-response amplitudes and all individual resting-state network amplitudes revealed that the relationship was limited to positive correlations with specific sensory and motor networks, and thus unlikely driven by potentially undesirable global signal properties (Supplementary Information: *Common fMRI global signal confound assessment*).

### Noxious-response amplitudes were associated with underlying white matter mean diffusivity

The SVR prediction model was trained on all infants in the nociception-paradigm dataset (n=18) to map from confound-adjusted resting-state network amplitudes to confound-adjusted noxious-response amplitudes. Using this model, predicted noxious-response amplitudes were generated for a 215-infant age-matched sample from the dHCP dataset using an identical approach for extracting resting-state network amplitudes and imaging confounds. The distribution of predicted noxious-response amplitudes in the dHCP dataset closely matched the distribution of the observed noxious-response amplitudes in the nociception-paradigm dataset (Figure 4 grey and blue histograms). These predicted noxious-response amplitudes were used for the structure-function analysis exploratory arm due to the large sample size. Findings were subsequently confirmed in the smaller nociception-paradigm dataset, which has true noxious-response amplitudes, constituting the structure-function analysis confirmatory arm.

**Figure 4:**
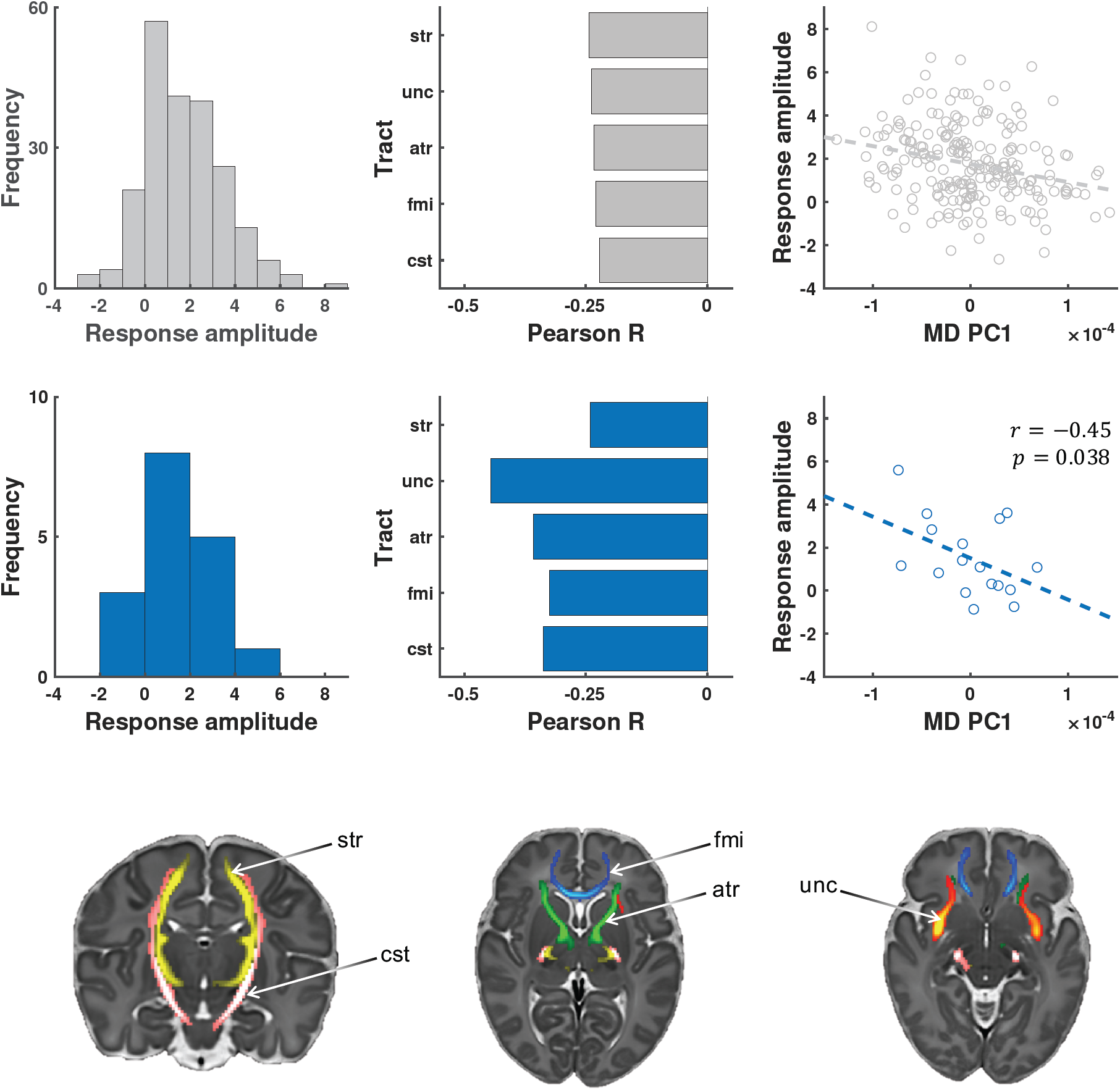
The relationship between noxious-response amplitude and white matter mean diffusivity. Top row: These three plots in grey (histogram, bar plot, scatter plot) are using the 215-infant dHCP dataset. Middle row: These three plots in blue are using the 17-infant nociception-paradigm dataset. Bottom row: These three maps display the five bilateral white matter tracts identified in the structure-function analysis exploratory arm (see Supplementary Figure 7). The histograms display the frequency distributions of the noxious-response amplitudes – predicted responses in dHCP dataset and observed responses in nociception-paradigm dataset. The bar plots display the Pearson correlation coefficients between noxious-response amplitudes and MD for the five white matter tracts. The scatter plots display the negative correlation between noxious-response amplitudes (y-axis) and MD PC 1. Abbreviations: atr = anterior thalamic radiation; cst = corticospinal tract; fmi = forceps minor; str = superior thalamic radiation; unc = uncinate fasciculus.

In the dHCP dataset, we performed an exploratory analysis to assess three dMRI microstructural parameters (mean diffusivity, fractional anisotropy, and mean kurtosis) across 16 bilateral white matter tracts. We found that the predicted noxious-response amplitudes were statistically significantly negatively correlated with mean diffusivity (MD) in five white matter tracts: anterior thalamic radiation (atr), corticospinal tract (cst), forceps minor (fmi), superior thalamic radiation (str), and uncinate fasciculus (unc) (Figure 4 grey bar plot and three maps; Supplementary Information: *Univariate correlations between noxious-response amplitudes and dMRI features*; Supplementary Figure 7). The first principal component of MD across these five tracts (MD PC1) accounted for 84.5% of the cross-infant variance, and as expected, was negatively correlated (r = –0.25) with the predicted noxious-response amplitudes (Figure 4 grey scatter plot). In summary, we used the large dHCP exploratory dataset to identify a network of 5 white matter tracts that have specific microstructural properties (characterised by their MD) that relate to the infants’ predicted amplitudes of noxious-evoked brain activity.

To validate these exploratory findings, we tested whether the actual noxious-evoked brain activity recorded in the infants in the nociception-paradigm dataset was also dependent on the same structural brain properties. We found that for each of the 5 white matter tracts the negative correlation coefficients between the MD of these tracts and the noxious-response amplitudes were present (Figure 4 blue bar plot). Additionally, MD PC1 accounted for 88.28% of the between-infant variance, and was statistically significantly negatively correlated with infants’ noxious-response amplitudes: r = –0.45, p-value = 0.038 (Figure 4 blue scatter plot). Thus, within our nociception-paradigm dataset, 20% (Pearson r^2^) of the between-infant variation in noxious-response amplitudes could be explained by the mean diffusivity of these five specific white matter tracts.

## Discussion

This study demonstrates that individual variability in newborn infant pain-related brain activity is dependent on the structural and functional architectures of the brain. By applying a mild experimental noxious stimulus to the infant’s foot, we quantified pain-related changes in brain activity, which are known to be similarly evoked by a range of tissue-damaging medical procedures, such as blood sampling, vaccinations, and cannulations ^3,19,20^. The tightly-controlled stimulus used in this study does not cause behavioural distress ^3^, but activates Aδ- fibres in the periphery and elicits noxious-evoked brain activity in the cerebral cortex ^15,21,22^, making it a useful experimental tool to better understand infant pain.

We took a multimodal MRI approach, using resting-state fMRI and white matter dMRI to understand between-subject differences in the amplitude of an infant’s cerebral response to a nociceptive input. We have shown that noxious-evoked brain activity is tightly coupled to resting-state network activity, and that the strength of this coupling is sufficiently robust to drive out-of-sample predictions. By observing a structure-function relationship between noxious-response activity and white matter microstructure, replicable in two independent datasets, we show that the infants’ observed noxious-response amplitudes also reflect specific stable trait effects, such as white matter myelination and fibre packing density. The ability to predict an infant’s trait cerebral haemodynamic response to nociceptive input from their resting-state brain activity highlights the potential use of infant resting-state brain activity to inform decision-making regarding pain management strategies for newborn infants.

Interpreting the newborn infant cerebral haemodynamic response amplitude to nociceptive input is challenging due to the lack of verbal report of the infants’ subjective experience. Here, the flexibility of MRI has allowed the identification of several novel neural correlates of the noxious-response amplitudes. In this study we observe a positive correlation between the amplitude of the noxious-response and the amplitude of resting-state network activity in sensory and motor networks in a group of healthy term-aged infants (Supplementary Figure 4). Given that the developmental trend from infancy to adulthood (observed using fMRI) is increasing haemodynamic response amplitude ^23^, this may suggest that the higher amplitude responses seen in some infants may reflect their increased structural and functional brain maturity. A number of published studies support this hypothesis. Using near-infrared spectroscopy to measure haemodynamic responses to pain in the perinatal period, the amplitude of these responses was also observed to progressively increase with age ^15^. And studies using fMRI to observe the developmental progression of resting-state activity have also found increased sensory and motor (and cognitive) network functional connectivity strength and activity amplitude with increasing age ^24,25^. While the structural correlates tested in our nociception-paradigm infant dataset demonstrated statistically significant negative correlation between noxious response amplitude and white matter MD, clear trends existed for both MD and FA. In both the nociception-paradigm and dHCP datasets, infants with larger noxious-response amplitudes (or for the dHCP data, larger predicted values) had smaller MD and larger FA values throughout the brain (Supplementary Figure 7). White matter MD decreases and FA increases throughout development into adulthood ^26^, and similar to the functional measures, these developmental trends are discernible within the perinatal period ^27^. The combination of negative MD and positive FA correlations suggest specific structural maturational influences, such as increasing white matter myelination, or fibre packing density, or both.

Taken together, the functional resting-state and structural white matter correlates suggest the infant noxious-response amplitude is a reflection of brain maturity, with larger response amplitudes indicating a more mature brain and negative and negligible response amplitudes indicating a more immature brain. This neural maturity hypothesis is consistent with the concept that infants’ noxious-response amplitudes are maturity dependent trait effects due to their dependency on underlying microstructure. This also suggests a plausible explanation as to why the infant’s resting-state activity can be used to predict the amplitude of the infants noxious-evoked brain response: both the activity levels recorded at rest and in response to stimulation are a function of each infant’s cerebral maturity and thus closely reflect each other due to a common underlying cause. Assuming increasing age is a reasonable proxy for increasing maturity, one can view an infant’s age as an imperfect indicator of the individual infant’s neural maturational state. Nevertheless, it is clear that two perfectly age-matched infants would not be expected to be perfectly matched for maturational state. The results of the current study suggest that a specific subset of MRI-measurable features allow us to detect individual variability in the maturity of the structural and functional neural architecture of the infant brain, which is not fully captured by alternative proxy indicators of neural maturity, including age and brain volume (Supplementary Figure 6). How well this hypothesis generalises to infants outside of the studied age-range or to non-healthy non-normative populations, such as infants born very prematurely then studied at ages 36-42 weeks PMA, would be a highly informative route of enquiry.

Further understanding of the biological interpretability of the noxious-response features will be required to appreciate the neurophysiological basis for this observed individual variability. The mild experimental stimulus used in this study likely evokes a multidimensional response profile in the infant brain including sensory discriminative aspects such as sharpness localised to the foot, cognitive aspects such as salience and attention, motor aspects such as post-stimulus movement, and potentially emotional aspects such as mild negative emotional valence. Disambiguating which aspect of the cerebral response is predictable from resting-state activity will be a challenge due to the limited behavioural repertoire of infants. However, it is possible to develop a principled approach for noxious-response feature extraction that could decrease this ambiguity and improve biological interpretability. There are now several candidate fMRI neural signatures for distinct components of adult pain and negative affect that could be applied to infant noxious-response data ^28–32^, and this is currently being actively researched by our group. Additionally, the reported functional coupling between infant resting-state and stimulus-response activities is currently limited to the nociceptive stimulus modality employed in this study. In adults, this functional coupling has been demonstrated for a wide range of tasks ^7,10,33^, and we imagine a similar generalisability of resting-state coupling to stimulus responses would be possible in newborn infants. While “task” fMRI experimental designs are severely limited in newborn infants, previous studies using non-nociceptive stimuli, such as non-noxious touch ^34^, auditory ^35^, and visual ^36^ stimuli have demonstrated the feasibility of a multimodal experimental design to test this directly. Finally, the functional coupling results may not generalise to premature infants younger than 35.9 weeks PMA, the youngest infant included in the present cohort. These younger infants will have less mature structural and functional brain architectures and poorer neurovascular coupling ^37,38^, which would need to be considered when investigating functional coupling.

A noteworthy feature of this study is the use of a larger publicly available multimodal dataset to enhance the findings within our smaller specialised nociception-paradigm dataset. This approach helps address three major challenges inherent to infant fMRI pain studies: limited sample size, limited literature base, and temporal stability. While these issues are not unique to infant fMRI pain studies, they are particularly challenging because of the combination of population and paradigm. First, small sample sizes are known to result in highly variable and unreliable accuracy in cross-validated prediction analyses ^39,40^. To validate the accuracy of our small sample size prediction model, we applied it to a sample of the dHCP dataset. Using predicted pinprick responses, we identified novel structure-function relationships that were subsequently confirmed in our nociception-paradigm dataset, underscoring the accuracy and meaningfulness of our prediction model’s outputs. Second, the limited literature base in newborn infant fMRI is highly problematic if researchers wish to engage in non-exploratory hypothesis-driven research. To date, there are only five newborn infant fMRI studies using nociception paradigms ^21,41–44^, two of which are technical papers looking at approaches to fMRI data acquisition ^43^ and analysis ^41^. To overcome the limited knowledge base in which we can formulate well-defined hypotheses to understand the noxious-related structure-function MRI associations in newborn infants, we used an exploratory-confirmatory analysis approach. Exploratory analyses were performed across a wide range of white matter tracts and diffusion parameters in the larger sample dHCP dataset in order to identify candidate associations, which were subsequently directly tested and confirmed in the smaller sample nociception-paradigm dataset. This two-armed approach allowed us to formulate data-driven hypotheses that could subsequently be empirically confirmed without double-dipping. Third, directly establishing the temporal stability of infants’ haemodynamic response amplitude to nociceptive input would involve multiple within-subject recordings, which is often not a viable approach. While there are studies in adults ^9,45^ demonstrating the temporal stability of static resting-state functional connectivity metrics (given reasonable data quality), analogous studies in neonates do not exist. We tested for temporal stability of noxious-response amplitudes through association with white matter microstructure, which is insensitive to wakefulness and physiological stress states, but highly sensitive to the integrity of developing structural connectivity, which constrains infants’ noxious-evoked responses. Using the large sample size dHCP dataset was, again, central to identifying the structure-function association in our nociception-paradigm dataset that accounted for 20% of the total cross-infant variance in noxious-response amplitudes. We believe the close association with white matter microstructure, coupled with the predictability from resting-state features, strongly suggests the infants’ noxious-response amplitudes are stable trait features of the brain.

Using multimodal MRI analyses, we have established that individual variability in pain-related brain activity in healthy peri-term-aged infants is tightly coupled to both the infants’ spontaneous resting-state activity and underlying white matter microstructure. Importantly, the amplitude of an individual infant’s noxious-response brain activity can be predicted from their spontaneous noxious-free resting-state brain activity. Even healthy newborn infants, within the first few days of postnatal life, display a wide range of responses to nociceptive input, likely a result of both genetic and environmental influences. This normative variability may reflect differences in individual resilience and vulnerability to environmental insults, such as clinical painful procedures that are frequently performed in hospitalised infants. The ability to predict an infant’s responses to pain and nociceptive input may have the potential to advance neonatal personalised pre-emptive pain management, and this study highlights the importance of understanding resting-state brain activity in achieving this goal. A better understanding of how individual differences in brain architecture influence pain processing is of paramount importance if we are to identify infants at increased risk of long-term alterations in brain structure and function and cognitive performance as a result of early life pain exposure. Early life pain and stress have the potential to alter an infant’s developmental trajectory and to influence their childhood well-being ^46,47^, but it may also increase the risk of developing chronic diseases in later life ^48,49^. The development of brain-based correlates of pain sensitivity could help identify vulnerable infants with the aim of tailoring pain relief treatments in a more principled, personalised, and evidence-based manner.

## Methods

### Part 1: Relating noxious-response amplitude to resting-state activity

#### Subject information

We recruited healthy neonates from the postnatal ward at the John Radcliffe Hospital (Oxford University Hospital NHS Trust). Infants were considered healthy if they were inpatients on the postnatal ward that never required admission to the neonatal unit, had no history of congenital conditions or neurological problems, and were clinically stable at the time of study. Written informed consent was obtained from parents prior to the study. Ethical approval was obtained from an NHS Research Ethics Committee (National Research Ethics Service, REC reference: 12/SC/0447), and research was conducted in accordance with standards set by Good Clinical Practice guidelines and the Declaration of Helsinki. Demographic details of the 18-infant sample are displayed in Table 2. Definitions of the age and total brain volume variables are detailed below (see *Clinical variables*).

**Table 2:** Demographic details of the 18-infant nociception-paradigm dataset. PMA = postmenstrual age; GA = gestational age; PNA = postnatal age; BW = birth weight; TBV = total brain volume; μ = mean; σ = standard deviation. * = excluded from dMRI analysis due to incomplete dMRI data.

| Infant | PMA<br>(weeks) | GA<br>(weeks) | PNA<br>(days) | BW<br>(grams) | TBV<br>(mm<sup>3</sup>) | Sex |
| --- | --- | --- | --- | --- | --- | --- |
| 1 | 40.6 | 40.3 | 2 | 3,880 | 283,685 | M |
| 2 | 37.4 | 37.1 | 2 | 4,570 | 276,222 | M |
| 3 | 35.9 | 35.3 | 4 | 1,910 | 212,410 | F |
| 4 | 35.9 | 35.6 | 2 | 3,180 | 284,623 | M |
| 5 | 38.3 | 38 | 2 | 3,400 | 309,632 | M |
| 6 | 38 | 37.3 | 4 | 3,410 | 306,530 | F |
| 7 | 39.6 | 39.3 | 2 | 3,250 | 301,086 | F |
| 8 | 36.4 | 36 | 3 | 3,510 | 260,268 | F |
| 9 | 38 | 37.9 | 1 | 2,490 | 221,792 | M |
| 10 | 40.7 | 40.6 | 1 | 4,300 | 346,861 | M |
| 11 | 40.4 | 40.1 | 2 | 4,040 | 356,454 | M |
| 12 | 39.3 | 39 | 2 | 3,775 | 300,094 | F |
| 13 | 38.9 | 38.6 | 2 | 2,950 | 286,714 | F |
| 14* | 41.7 | 41.4 | 2 | 3,400 | 297,004 | M |
| 15 | 40.4 | 39 | 10 | 3,750 | 416,904 | F |
| 16 | 38.3 | 38 | 2 | 2,780 | 247,199 | M |
| 17 | 37.4 | 36.4 | 7 | 2,235 | 257,069 | F |
| 18 | 39 | 38.9 | 1 | 3,350 | 252,271 | M |
| $\mu$ | 38.7 | 38.3 | 2.8 | 3,343.3 | 289,823.2 | --- |
| $\sigma$ | 1.7 | 1.8 | 2.3 | 690.4 | 49,020.8 | --- |

#### Experimental setup and design

Neonates were transported to the Wellcome Centre for Integrative Neuroimaging (Oxford, UK), then fed and swaddled prior to scanning. Infants were fitted with ear plugs, ear-muffs, and ear-defenders, and placed on a vacuum-positioning mattress with additional soft padding around the head to restrict motion. Heart rate and blood oxygen saturation were monitored throughout scanning. An event-related experimental design was used for the nociception paradigm ^21^. The mild non-skin-breaking nociceptive stimulus was a 128 mN sharp-touch pinprick (PinPrick Stimulator, MRC Systems). Ten trials of the stimulus were delivered to the dorsum of the left foot, each trial was 1 s, and the minimum inter-stimulus interval was 25 s. This long inter-stimulus interval was used to minimise the influence of motion at the time of stimulus delivery. The stimuli were applied when the infants were naturally still. For all other scan types, infants lay passively in the scanner. No sedatives were used at any stage of this study.

#### MRI data acquisition

All data were collected on a 3T Siemens Prisma with an adult 32 channel receive coil. The structural data acquisition was: T2-weighted, TSE (factor 11), 150° flip angle, TE = 89 ms, TR = 14,740 ms, parallel imaging GRAPPA 3, 192 × 192 in-plane matrix size, 126 slices, 1 mm isotropic voxels, and 2 mins 13 s acquisition time. The fieldmap data acquisition was: gradient echo, 2DFT readout, dual echo TE1/TE2 = 4.92/7.38 ms, TR = 550 ms, 46° flip angle, 90 × 90 in-plane matrix size, 56 slices, 2 mm isotropic voxels, and 1 min 40 s acquisition time. Both the resting-state and noxious-response fMRI data acquisitions were: T2* BOLD-weighted, gradient echo, EPI readout, 70° flip angle, TE = 50 ms ^43^, TR = 1,300 ms, multiband 4 ^50,51^, 90 × 90 in-plane matrix size, 56 slices, 2 mm isotropic voxels, AP phase encode direction, and a single-band reference (SBref) image was acquired at the start. Resting-state acquisition time was 10 mins 50 s (500 volumes), and noxious-response mean acquisition time was approximately 6 min (approximately 277 volumes). The dMRI data acquisition was: T2 diffusion-weighted, spin echo, EPI readout, 90° flip angle, TE = 73 ms, TR = 2,900 ms, multiband 3, 102 × 102 in-plane matrix size, 60 slices, 1.75 mm isotropic voxels, AP phase encode direction, multishell (b = 500, 1000, 2000 s/mm^2^), a total of 140 directions uniformly distributed over the whole sphere, and approximately 8 mins acquisition time. Phase-reversed b0 images were collected to derived a spin-echo fieldmap for distortion correction of the diffusion data.

#### MRI data preprocessing

All MRI data were preprocessed using analysis pipelines developed as part of the Developing Human Connectome Project (dHCP) (http://www.developingconnectome.org). The T2 structural data were processed (brain extraction, bias field correction, and tissue segmentation) using the MIRTK Draw-EM neonatal pipeline ^52^, the tool forming the basis of the dHCP structural preprocessing pipeline ^53^. The GRE dual-echo fieldmap data were processed using a modified version of fsl_prepare_fieldmap.

Both the noxious-response and resting-state fMRI data were preprocessed using an extended version of the dHCP fMRI preprocessing pipeline ^24,41^. The functional data were corrected for motion and distortion using FSL’s EDDY ^54,55^, which included slice-to-volume motion correction ^56^ and susceptibility-by-movement distortion correction ^57^. Noxious-response fMRI data were high-pass temporally filtered at 0.01 Hz, and resting-state fMRI data at 0.005 Hz. Data were then denoised using FSL’s FIX ^58,59^, low-pass spatially filtered with a 3 mm FWHM filter using FSL’s SUSAN ^60^, and grand mean scaled to a global spatiotemporal median of 10,000. For spatial normalisation to standard space ^61^, the data were first registered from functional space to the infant’s T2 structural space, via the SBref, using 6 DoF rigid-body alignment, refined using BBR ^62^ with FSL’s FLIRT ^63,64^. The registration from structural space to the 40-week template ^61^, via an age-matched standard template, was performed using ANTs’s SyN ^65^.

The diffusion data were analysed using the dHCP dMRI preprocessing pipeline ^27,66^. The blip-up and blip-down b0 images were used to generate the fieldmap using FSL’s TOPUP ^67,68^. The diffusion data were simultaneously corrected for motion, distortion, and eddy currents using FSL’s EDDY, which included outlier detection and replacement ^69^ as well as the slice-to-volume motion correction and susceptibility-by-movement distortion correction used in the fMRI data correction. Spatial normalisation followed the same sequence of registrations as the functional data.

#### Noxious-response amplitudes

For each infant, a noxious-response map was generated using standard subject-level voxelwise GLM analysis in FSL’s FEAT ^70^, fitting the term-neonate double-gamma HRF ^23,41^. A group average t-statistic map was generated using the 18 infants’ noxious-response regression parameter maps. Individual infants’ regression parameter maps were used as the subject-level noxious-response maps; the group-average t-statistic map was used as the group-level noxious-response map. To summarise an individual infant’s noxious-response map to a single scalar measure of noxious-response amplitude, the group-level noxious-response map was regressed onto the infant’s noxious-response map. Thus, an infant’s noxious-response amplitude was defined as this spatial regression coefficient. Assessment of the potential influence of HRF goodness-of-fit on noxious-response amplitudes is detailed in Supplementary Information (*Noxious-response HRF fit assessment* and Supplementary Figure 1,3)

As detailed below, our prediction analyses examined associations between these noxious-response amplitudes and three sets of predictors (resting-state network amplitudes, resting-state imaging confounds, and clinical variables), and our structure-function analyses examined associations between noxious-response amplitudes and a dMRI model parameter (mean diffusivity). In all these analyses, the noxious-response amplitudes were adjusted for a set of three noxious-response imaging confounds extracted from each infant’s noxious-response fMRI data: mean head motion, stimulus-correlated head motion, and CSF signal amplitude. These three metrics were intended to capture cross-infant noxious-response variability due to subject motion (mean and stimulus-correlated head motion) and cardiac pulsatility (CSF amplitude). Mean head motion was defined as the mean framewise displacement across the entire noxious-response fMRI scan session. Stimulus-correlated head motion was estimated as a multiple correlation coefficient z-statistic (Fisher r-to-z transformation) between the predicted BOLD response (stimulus application timeseries convolved with the HRF) and the 24 head motion parameter timeseries (estimated by EDDY during motion correction). CSF amplitude was estimated from each infant’s noxious-response map as the mean regression coefficient within the CSF ROI. Details on the CSF ROI construction are provided in Supplementary Information (*CSF and white matter regions-of-interest definition* and Supplementary Figure 9).

#### Resting-state network amplitudes

To define a robust set of core resting-state networks in the infants’ resting-state fMRI data, resting-state networks identified in the 18-infant nociception-paradigm dataset were compared to those identified in a subset of the dHCP dataset, which had previously been produced as part of the dHCP ^14^. A robust set of core resting-state networks was defined as those replicated across datasets. Demonstrating replicability in the dHCP dataset confirmed the set of core networks were robust and not unique to our nociception-paradigm dataset. The dHCP data subset included 242 healthy term-aged infants: mean GA at birth = 38.6 weeks; mean PMA at scan = 40.4 weeks; 112 females and 124 males. The resting-state network analysis performed on the 18-infant nociception-paradigm dataset was closely matched to that described for the dHCP dataset ^24^. In brief, probabilistic functional mode (PFM) analysis using FSL’s PROFUMO ^17,18^ was run on both datasets with a pre-specified dimensionality of 25, and using the infant double-gamma HRF ^23,41^ as the temporal prior. PROFUMO’s Bayesian model complexity penalties can eliminate modes, thus returning a number of group-level modes that can be less than the pre-specified dimensionality. This is noted, as the data-determined dimensionality of the nociception-paradigm dataset was 11 despite a pre-specified dimensionality of 25. Due to the larger sample size, the data-determined dimensionality of the dHCP dataset was equal to the pre-specified dimensionality. The dHCP resting-state network maps had greater SNR due to the significantly larger sample size. Thus, the dHCP resting-state network maps forming the set of core resting-state networks were used as the template maps to extract resting-state network amplitudes from the resting-state fMRI data.

These resting-state network template maps were spatially regressed onto each infant’s resting-state functional data using multiple regression, resulting in network timeseries. While the timeseries standard deviation is the typical amplitude metric used and is the default in FSL’s FSLNets ^71^, the standard deviation is sensitive to outliers, which in this context, typically appear as head motion-related timeseries spikes. Resting-state network amplitudes were thus quantified using the median absolute deviation (MAD), due to the MAD’s increased robustness to outliers. This set of resting-state network amplitudes was directly tested for association with noxious-response amplitudes after adjusting for resting-state imaging confounds (defined below). Assessment of the potential influence of resting-state network timeseries outliers on noxious-response amplitudes is detailed in Supplementary Information (*Resting-state network timeseries outlier assessment* and Supplementary Figure 2,3).

#### Resting-state imaging confounds

A set of resting-state imaging confounds was directly tested for association with noxious-response amplitudes and used for confound-adjusting the resting-state network amplitudes. These confounds included three metrics extracted from each infant’s resting-state fMRI data: mean head motion, CSF amplitude, and white matter amplitude. These three metrics were intended to capture cross-infant resting-state variability due to subject motion (mean head motion), cardiac pulsatility (CSF amplitude), and global signal (white matter amplitude). Directly testing these resting-state imaging confounds assessed whether associations between resting-state network amplitudes and noxious-response amplitudes could be explained by undesirable artefactual features of the resting-state data. Mean head motion was defined as the mean framewise displacement across the entire resting-state fMRI scan session. Mean CSF and white matter timeseries were extracted from each infant’s resting-state data, and the timeseries amplitudes were defined as the MAD of these timeseries. Details on the CSF and white matter ROI construction are provided in Supplementary Information (*CSF and white matter regions-of-interest definition* and Supplementary Figure 9).

#### Clinical variables

A set of clinical variables was directly tested for association with noxious-response amplitudes and included the six variables in Table 2: postmenstrual age (PMA), gestational age (GA), postnatal age (PNA, also called chronological age), birth weight (BW), total brain volume (TBV), and sex. The three age variables are defined according to the American Academy of Paediatrics ^72^, and the TBV was calculated from the infants’ structural MRI data using the tissue segmentation outputs of the structural preprocessing pipeline. Testing the clinical variables assessed whether associations between resting-state network amplitudes and noxious-response amplitudes could be explained by biologically interesting underlying variables.

#### Predicting noxious-response amplitudes

For all prediction analyses, the responses to be predicted were the infants’ whole-brain noxious-response amplitudes (Figure 1 scalar values). Three sets of predictors were tested for predictive capacity: nine resting-state network amplitudes, six clinical variables, and three resting-state imaging confounds. For all three sets of predictors, a support vector regression (SVR) model with a linear kernel was used. Linear SVR was selected over a linear regression via ordinary least squares, due to the SVR cost function’s greater robustness to outliers. Out-of-sample predictions were generated using leave-one-out cross validation (LOO-CV). The noxious-response amplitudes were confound-adjusted for the three noxious-response imaging confounds (mean head motion, stimulus-correlated head motion, and CSF amplitude). When generating predictions using the resting-state network amplitudes, this set of predictors was confound-adjusted for the three resting-state imaging confounds (mean head motion, CSF amplitude, and white matter amplitude).

The linear SVR model was fit in Python using scikit-learn packages ^73^, with all steps performed in a LOO-CV manner. Confound adjustment of the resting-state network amplitudes and noxious-response amplitudes was performed using cross-validated confound regression, implemented using the publicly available code by Lukas Snoek (https://github.com/lukassnoek/MVCA), as described in the author’s article ^74^. Responses were z-scaled using training set means and standard deviations. The scikit-learn SVR parameters were: kernel = linear, loss function = epsilon insensitive, epsilon = 0.1, regularization = ridge, regularization strength = {0.001, 0.01, 0.1, 1}. Optimisation of the regularization strength parameter was performed using an initial LOO-CV grid search over this set of values. Regularisation tuning and SVR model training were optimised to minimise mean squared error.

The prediction accuracy was assessed using three summary metrics: root mean squared error (RMSE), sums-of-squares formulation of the coefficient of determination (R^2^), and Spearman’s rank correlation coefficient (R_Sp_). The RMSE was selected as the primary metric of prediction accuracy, as it directly quantifies the error (the difference between predicted and actual observed values) and is in original units. The R^2^ was also reported, as its value is interpreted as the proportion of the total variation of the response (about its mean) that is accounted for by the fitted model, and is thus an intuitive metric to assess success of the predictions. The R_Sp_ between predicted and observed noxious-response amplitudes was also reported, as it may be valuable to know the model’s ability to correctly rank infants’ noxious-response amplitudes, on a relative scale, from lowest to highest. To test the statistical significance of the RMSE, R^2^, and R_Sp_ measures using null hypothesis testing, one-tailed significance tests were performed using permutation analysis, running 1,000 permutations through the full prediction pipeline. Assessment of the potential influence of an fMRI global signal confound (common to both resting-state and noxious-response data) on noxious-response amplitudes is detailed in Supplementary Information (*Common fMRI global signal confound assessment* and Supplementary Figure 4)

### Part 2: Relating noxious-response amplitude to white matter microstructure

#### Structure-function analysis using an exploratory-confirmatory approach

The infants’ noxious-response amplitudes were assessed for structure-function associations by analysing white matter microstructure to better understand the biological basis for individual variability in noxious-response amplitude and to evaluate the temporal stability of the observed responses. Due to the insensitivity of white matter microstructure to wakefulness and emotional state, an observed structure-function relationship would suggest the infants’ noxious-response amplitudes were temporally stable trait effects. Temporal stability was assessed using this structure-function approach rather than looking at stability across multiple test occasions, as infants could only be tested on a single occasion. Due to the lack of knowledge regarding the brain’s structural basis for noxious responses in healthy newborn infants, an exploratory analysis was required. However, due to the small sample size of the nociception-paradigm dataset (n=17 with dMRI data, Table 2), the appropriate statistical multiple testing corrections to control the false positive rate would prohibit the identification of a true positive. To overcome this issue, we adopted an exploratory-confirmatory analysis approach. We used a large age-matched sample from the dHCP dataset (n=215, sample defined below) for the exploratory arm, in which a wide range of white matter tracts and dMRI model parameters were studied in order to identify candidate nociception-relevant microstructural features. Structure-function relationships identified in this exploratory arm facilitated the formulation of specific well-defined hypotheses. These were subsequently tested in the nociception-paradigm dataset (n=17) for validation, which constituted the confirmatory arm of the analysis.

#### Noxious-response amplitudes in the dHCP dataset

The dHCP fMRI data includes resting-state data only. To analyse nociception-relevant structure-function relationships in this dataset, the dHCP resting-state data were mapped to noxious-response amplitudes using the SVR prediction model described previously – see *Predicting noxious-response amplitudes* above. This prediction model was trained on the nociception-paradigm dataset (n=18) using the nine resting-state network amplitudes as predictors (adjusted for resting-state imaging confounds) and the noxious-response amplitudes as responses (adjusted for noxious-response imaging confounds). In a sample from the dHCP dataset (defined below), the nine resting-state network amplitudes and three resting-state imaging confounds were extracted in an identical manner to the analysis performed in the nociception-paradigm dataset – see *Resting-state network amplitudes* and *Resting-state imaging confounds* above. The resting-state network amplitudes were adjusted for the resting-state imaging confounds, and the adjusted amplitudes were used to generate predicted noxious-response amplitudes. Frequency distribution histograms of the predicted noxious-response amplitudes from the dHCP dataset and the observed noxious-response amplitudes from the nociception-paradigm dataset were qualitatively compared.

#### Sample selection in the dHCP dataset

Infants in the dHCP dataset were included in our sample if they satisfied three quality control (QC) criteria and two age criteria to ensure the sample data were of reasonable quality and were age-matched to the prediction model training set. The three QC criteria were: (i) both an infant’s fMRI and dMRI data had to pass basic dHCP QC pipelines ^24,66^, (ii) both scan sessions had to have completed fully (300 volumes for dMRI data; 2,300 volumes for fMRI data) to remove inter-subject variability due to data quantity related to scan length (all infants in the nociception-paradigm dataset satisfy this criterion), and (iii) the vertex of the cerebral cortex had to remain within the scan field of view (FOV) for at least 95% of scan session (all infants in the nociception-paradigm dataset satisfy this criterion). This last QC criterion excluded infants in which primary somatosensory and motor brain regions, demonstrated in the nociception-paradigm dataset to be of central importance to noxious stimulus processing (Supplementary Figure 4), would have unreliable data. The two age criteria were: (i) infants had to have both a gestational age and a postmenstrual age at time of scan between 36–42 weeks, and (ii) infants had to have been scanned within the first 10 days of postnatal life. These selection criteria resulted in a dHCP dataset sample size of n=215 infants.

#### White matter microstructural features

Analogous to the pre-existing dHCP resting-state network templates used for resting-state network amplitude feature extraction (see *Resting-state network amplitudes* above), our 215- infant dHCP sample was used to generate a set of 16 bilateral white matter tract regions-of-interest (ROIs). These tracts were generated using the “baby autoPtx” approach established as part of the dHCP dMRI preprocessing pipeline development ^27^. In brief, FSL’s probabilistic multi-shell ball and zeppelins model ^75^ is fit as part of the dHCP dMRI preprocessing pipeline. Probabilistic tractography using FSL’s PROBTRACKX ^76,77^ is run using pre-defined seed, target, and exclusion masks. At the time of analysis, masks for 29 white matter tracts were available, of which 13 were unilateral and three bilateral. To create bilateral white matter ROIs analogous to our bilateral resting-state networks, the unilateral tracts were fused, resulting in a total of 16 bilateral tracts. In our 215-infant dHCP sample, the normalised probability value results of each tract were group-averaged in standard space and thresholded at a probability of 0.01. As part of the dHCP preprocessing pipeline, FSL’s DTIFIT is used to generate mean diffusivity (MD), fractional anisotropy (FA), and mean kurtosis (MK) parameter maps for each infant. We thresholded each infant’s parameter maps to remove noisy voxels with values falling outside the expected theoretical range, which can happen in practice due to poor SNR or head motion: for MD, this included negative values; for FA, this included values outside the interval [0,1]; for MK, this included values outside the interval [0,3]. The 16 bilateral white matter ROIs were used to extract mean parameter values for each tract. These 48 values (16 tracts x 3 parameters) per subject constituted the white matter microstructural features for our structure-function analyses.

#### Identifying a valid structure-function association

Using the 215-infant dHCP sample, univariate correlations between predicted noxious-response amplitudes and each microstructural feature was assessed using permutation testing with FSL’s PALM ^78^. These correlations were adjusted for three dMRI imaging confounds: mean head motion (estimated by EDDY during preprocessing), number of noisy voxels falling outside the expected theoretical range (see *White matter microstructural features* above), and TBV (see *Clinical variables* above). Our dMRI parameters-of-interest are influenced by tissue density and partial voluming artefacts due to brain volume variance across infants, so adjustment for TBV was included to mitigate these global confounds. There is no need to adjust for fMRI imaging confounds, as the SVR prediction model maps to confound-adjusted noxious-response amplitudes. Statistical significance was assessed using two-tailed Pearson correlations with 10,000 permutations and FWER-corrected for multiple testing across all 48 tests ^79^. While the observed statistically significant negative correlations with the MD of five tracts (Figure 4 and Supplementary Figure 7) are statistically valid due to appropriate FWER-adjustment of false positive rate, these findings are tentative due to the use of predicted noxious-response amplitudes. The dHCP dataset has no noxious-response amplitude ground truth, so the results of this exploratory arm need confirmation in the nociception-paradigm dataset, for which ground truth observed noxious-response amplitudes exist.

In the confirmatory arm, the negative correlation between predicted noxious-response amplitudes and MD identified in the dHCP dataset was assessed in the nociception-paradigm dataset using two approaches. In both approaches, correlations were adjusted for both noxious-response imaging confounds (mean head motion, stimulus-correlated head motion, and CSF amplitude) and dMRI imaging confounds (mean head motion, number of noisy voxels, and TBV). First, the correlation polarities between observed noxious-response amplitudes and MD of the five statistically significant tracts were qualitatively compared across datasets. Thus, confirmatory arm question one was: “*are the correlation coefficient polarities (positive or negative) between noxious-response amplitudes and MD consistent between datasets for these five tracts?*” Second, in the dHCP dataset, principal component analysis was run across the MD values of the five tracts of all 215 infants. The first principal component (PC1) accounted for 84.5% of the total MD variance with a negative correlation with predicted noxious-response amplitudes (r = –0.25). Due to the double-dipping circularity of this analysis in the dHCP dataset ^80^, this negative correlation between MD PC1 and predicted noxious-response amplitude will be biased toward high statistical significance. However, the demonstration in the dHCP dataset that MD PC1 accounts for a major portion of the variance in these tracts and has a statistically significant negative correlation with predicted noxious-response amplitudes serves as a single straight-forward quantitative test that can be directly confirmed (or not) in the nociception-paradigm dataset in an unbiased and non-circular manner. Thus, confirmatory arm question two was: “*is the statistically significant negative correlation between noxious-response amplitudes and MD PC1 (across these five tracts) consistent between datasets?*”. Statistical significance was assessed in PALM using a one-tailed Pearson correlation with 10,000 permutations.

## Acknowledgements

We are grateful for the provision of simultaneous multi-slice (multi-band) pulse sequence and reconstruction algorithms from the Centre for Magnetic Resonance Research, University of Minnesota.

The dHCP resting-state network data were provided by the developing Human Connectome Project, KCL-Imperial-Oxford Consortium funded by the European Research Council under the European Union Seventh Framework Programme (FP/2007-2013) / ERC Grant Agreement no. [319456]. We are grateful to the families who generously supported this trial.

## Author Contributions

LB conceived the idea for the study, analysed the data, interpreted the results, and wrote the paper. FM collected the data and revised the paper. SF provided technical assistance with the dHCP fMRI preprocessing pipeline, provided the dHCP resting-state network maps, and revised the paper. MA analysed the data and revised the paper. RM analysed the data and revised the paper. MB provided technical assistance with the dHCP dMRI preprocessing pipeline, provided the dHCP white matter tract maps, and revised the paper. RR interpreted the results and revised the paper. SJ designed the experiment, interpreted the results, and revised the paper. ED designed the experiment, interpreted the results, and revised the paper. RS conceived the idea for the study, designed the experiment, interpreted the results, and revised the paper.

