## Supplementary information for "Functional and diffusion MRI reveal the functional and structural basis of infants’ noxious-evoked brain activity"

### Supplementary methods and results

##### Noxious-response HRF fit assessment

Voxels in each infant's noxious-response map were segmented into three classes using FSL's mixture modelling approach: null, positive BOLD, and negative BOLD [1]. Peristimulus timeseries plots were constructed using the default method in FSL's FEAT, using the noxious-response map voxel with largest z-statistic. Using the mixture modelling outputs, the voxel with largest positive z-statistic was extracted from the positive BOLD response class, and the voxel with largest negative z-statistic was extracted from the negative BOLD response class. This approach ensured the selected voxels do not come from regions lacking activation (i.e. the null class) but with a spurious good fit, and that the selected voxels are representative of the best HRF fit for that subject (i.e. the largest z-statistic). The peristimulus timeseries plots include all timepoints for all trials, the trial average response, and the HRF fit (Supplementary Figure 1).

To assess goodness-of-fit between the data trial average response and the HRF fit, the Pearson correlation coefficient was calculated and converted to a z-score using the Fisher r-to-z transformation. To assess the influence that alignment between the raw data and the fitted HRF may have on the infant's estimated noxious-response amplitudes, we performed a Pearson correlation test. Statistical significance was assessed in FSL's PALM [2] using a two-tailed permutation test with 10,000 permutations (Supplementary Figure 3 left). Due to the scale invariance of the Pearson correlation coefficient, the correlation should indicate the quality of alignment between the HRF and the data and not capture amplitude effects. There was no observed relationship between an infant's noxious-response amplitude and the quality of the HRF fit ($r = -0.075$, $p = 0.77$) suggesting that the cross-infant variability in noxious-response amplitudes is unlikely due to the degree of temporal misalignment when fitting the HRF.

##### Resting-state network timeseries outlier assessment

We quantified resting-state network amplitudes using the median absolute deviation (MAD) of the network timeseries. The MAD was selected over the standard deviation due to MAD's increased robustness to outliers. The presence of outliers in network timeseries indicate sudden changes in signal intensity, which can indicate head motion contamination. The underlying BOLD fluctuations should follow slow and smooth fluctuations, so the number of outliers across networks may give an indication of how faithfully our network timeseries reflect these BOLD fluctuations. Outliers were independently detected for each network timeseries using the "quartiles method": outliers defined as values more than 1.5 interquartile ranges above the upper quartile or below the lower quartile (Supplementary Figure 2).

For each infant, a count of total number of outliers across all nine networks was made. Larger outlier counts were taken to indicate a poorer relationship between the observed network timeseries and the underlying BOLD timeseries. While the presence of these outliers will not directly influence our measures of network amplitudes (due to the use of MAD), the number of outliers may indicate the quality of the network timeseries underpinning the estimated network amplitudes. To assess the potential relationship between an infant's noxious-response amplitude and the quality of the resting-state network timeseries, we performed a Pearson correlation test. Statistical significance was assessed in PALM using a two-tailed permutation test with 10,000 permutations (Supplementary Figure 3 middle). There was no observed relationship between an infant's noxious-response amplitude and the quality of the resting-state network timeseries ($r = -0.014$, $p = 0.95$) suggesting that our observation of functional coupling between noxious-response and resting-state activities is unlikely due to a resting-state network timeseries quality confound.

For completeness, we assessed the potential for a relationship between an infant's resting-state network timeseries quality and noxious-response HRF fit quality by performing a Pearson correlation test between infants' outlier counts (from resting-state data) and HRF z-scores (from noxious-response data) (Supplementary Figure 3 right). Statistical significance was assessed in PALM using a two-tailed permutation test with 10,000 permutations. The relationship was not statistically significant ($r = -0.47$, $p = 0.055$). There was a trend towards significance, but this was strongly influenced by an outlier. Interestingly, the outlier subject had the poorest HRF fit (Supplementary Figure 1 row 1 column 3), the poorest resting-state network timeseries quality (Supplementary Figure 2 row 1 column 3), and a negligible noxious-response amplitude (Figure 1 row 1 column 3). It is uncertain whether the low quality of this infant's resting-state and noxious-response features was due to artefactual low data quality or biologically interesting negligible BOLD signal. Future work examining this distinction will be vital to improving BOLD signal interpretation of the immature newborn infant's brain.

**Common fMRI global signal confound assessment**

The potential for an artefactual global signal present throughout the brain in both the resting-state and noxious-response runs could cause artefactual similarities between an infant's noxious-response and all resting-state network amplitudes. Univariate Pearson correlation analyses were performed between the noxious-response amplitudes and each individual resting-state network amplitude to test for global or network-specific associations (Supplementary Figure 4). Statistical significance was assessed in PALM using a two-tailed permutation test with 10,000 permutations. These results revealed that the relationship was not global, but network-specific. Infants' noxious-response amplitudes were significantly positively correlated with the resting-state network amplitudes of somatomotor and visual networks only (SMN and VNop in Figure 2): $r = 0.73$ ($p = 0.0003$) for SMN, and $r = 0.77$ ($p = 0.0001$) for VNop. This network specificity suggested that our observation of functional coupling between noxious-response and resting-state activities is unlikely due to a common global signal, which might have indicated an undesirable artefact of global signal properties.

Given the tactile nature of our nociception paradigm, the correlation between noxious-response amplitudes and resting-state SMN activity is unsurprising and may indicate a direct causal link. However, the correlation with resting-state VNop activity was unexpected. We do not believe the correlation is indicative of a direct causal link in a manner akin to the SMN relationship, but may instead be a non-causal association downstream of a deeper common cause. But why this potential underlying common cause would be limited to SMN and VNop is, again, an interesting question requiring further investigation.

For completeness, univariate Pearson correlations between noxious-response amplitudes and each of the resting-state imaging confounds and clinical variables are also reported (Supplementary Figure 5-6). In all cases, statistical significance was assessed in PALM using a two-tailed permutation test with 10,000 permutations, and no significant associations were found.

**Univariate correlations between noxious-response amplitudes and dMRI features**

In the structure-function analysis, we defined 16 bilateral white matter tract ROIs and assessed cross-infant variation in these tracts for three dMRI parameters: mean diffusivity (MD), fractional anisotropy (FA), and mean kurtosis (MK). In the 215-infant dHCP dataset, we correlated the infants' predicted noxious-response amplitudes with these 48 dMRI features (16 tracts x 3 parameters). The Pearson correlation coefficients for all features are displayed in Supplementary Figure 7 (grey bars). Five statistically significant negative correlations with MD were identified and are indicated with asterisks in Supplementary Figure 7 (highlighted with red box). Statistical significance was assessed in PALM using two-tailed permutation tests with 10,000 permutations and FWER-corrected for multiple testing across all 48 tests [3].

To qualitatively assess patterns in correlation coefficients consistent between the dHCP and nociception-paradigm datasets, the equivalent Pearson correlation coefficients were calculated in the nociception-paradigm dataset and plotted adjacent to the dHCP coefficients in Supplementary Figure 7. Testing for statistical significance in the nociception-paradigm dataset was not performed, as explained in the main text. While there does not seem to be any appreciable consistency between datasets for MK, a clear pattern emerges for both MD and FA. For MD, all 16 tracts exhibit negative correlations with noxious-response amplitudes in both datasets. For FA, all 16 tracts exhibit positive correlations with (predicted) noxious-response amplitudes in the dHCP dataset, and all bar one (medial lemniscus) exhibit positive correlations with (observed) noxious-response amplitudes in the nociception-paradigm dataset. Together, these results highlight very similar patterns of structure-function associations for both MD and FA across datasets, underscoring the sensibility of the resting-state-derived noxious-response amplitudes in the dHCP dataset. Additionally, the ubiquity of the negative MD correlations and positive FA correlations suggest that the nociception-related structure-function association is a global effect common to all white matter tracts to varying degrees. The lack of MK's consistency among tracts within individual datasets and within a tract between datasets may be due to a true lack of a structure-function association with this parameter, may be a failure of our prediction model to map from resting-state to noxious-response amplitudes with high enough accuracy for this parameter, or may simply be a data quality issue due to the known high noise levels in this parameter.

The five tracts identified in the dHCP dataset were confirmed using the nociception-paradigm dataset. Due to the small sample size (n=17), only a single hypothesis was tested in the nociception-paradigm dataset focusing on the first principle component across these five tracts (see Figure 4 in main text). For completeness, univariate correlations between noxious-response amplitudes and each of these tracts are displayed in Supplementary Figure 8.

**CSF and white matter regions-of-interest definition**

When associating noxious-response amplitudes with both resting-state fMRI features and white matter dMRI features, the functional data were adjusted for several imaging confounds. The noxious-response amplitudes were adjusted for three noxious-response imaging confounds (mean head motion, stimulus-correlated head motion, CSF amplitude), and the resting-state network amplitudes were adjusted for three resting-state imaging confounds (mean head motion, CSF amplitude, and white matter amplitude). The extraction of CSF and white matter amplitudes from the functional data required CSF and white matter ROI masks.

Due to the small size of the infant brain, partial volume contamination is problematic. The CSF and white matter masks used in our analyses were conservative to minimize grey matter contamination, and were defined using the dHCP neonatal template atlas [4]. The CSF ROI (Supplementary Figure 9 blue) is restricted predominantly to the fluid surrounding the brainstem and cerebellum and is a single region crossing the midline. Ventricular CSF was excluded from this ROI due to the small size of infant ventricles. The white matter ROI (Supplementary Figure 9 red) is restricted to the largest white matter regions in the cerebrum, consisting of a left and right half tightly localized to the centre of white matter regions. In the resting-state data, these ROIs were used to extract mean timeseries for both CSF and white matter. The amplitudes of these timeseries were quantified as the timeseries MAD values. In the noxious-response data, the CSF ROI was used to extract the mean regression parameter from infants' noxious-response maps, which constituted the noxious-response CSF amplitude.

### Supplementary figures

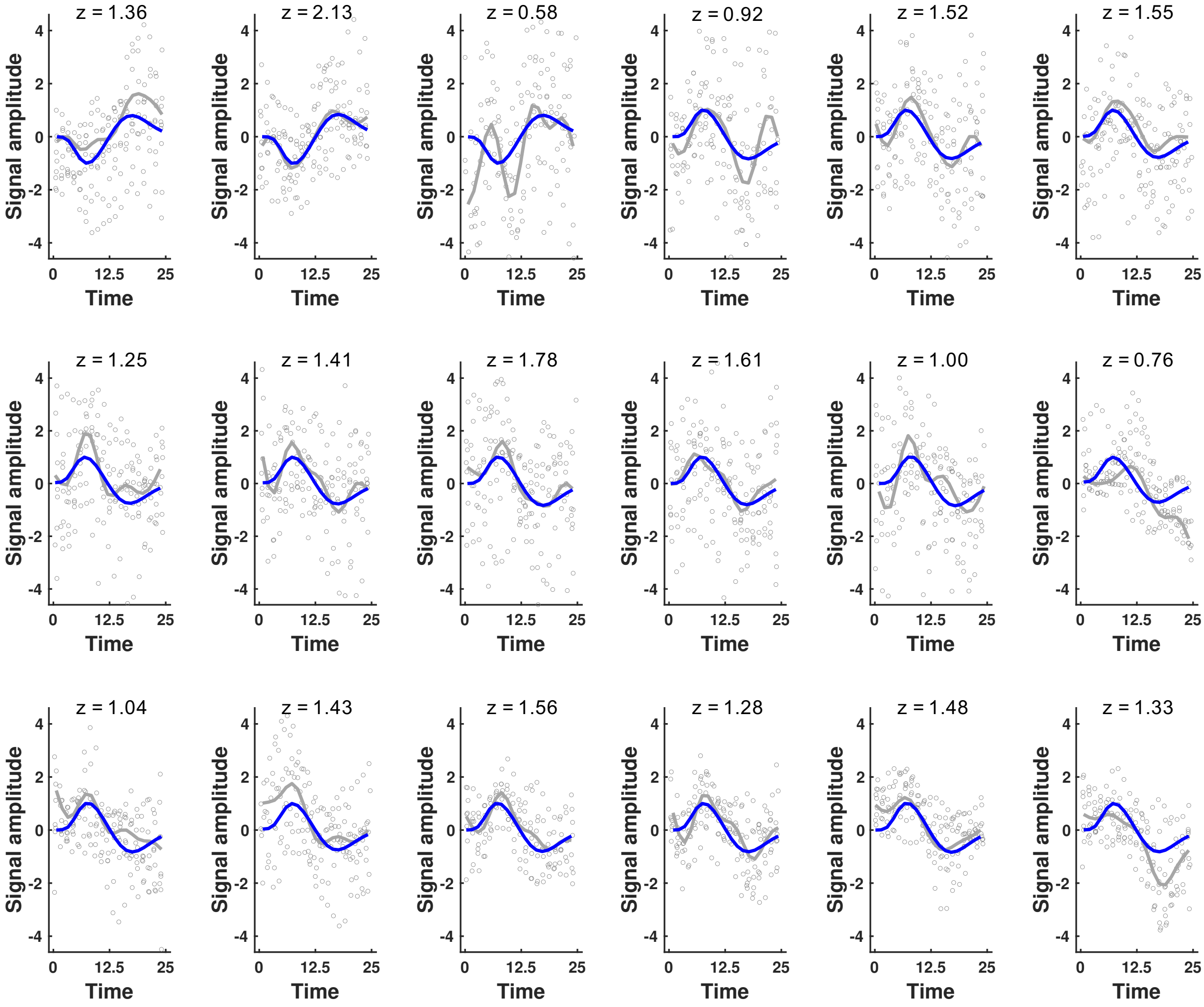


**Supplementary Figure 1: Noxious-response HRF fit.** The plots are displayed in identical order to Figure 1 (main text). For all plots, the x-axis (time in seconds) runs from 0 to 25, where t = 0 s is the point of stimulus delivery, and the minimum inter-stimulus interval is 25 s. The y-axis is arbitrarily scaled to blue curve (HRF fit) peak equals one. Grey circles are individual timepoints from all 10 trials, the grey line is the trial average timeseries, and the blue line is the HRF fit to the data. Above each plot is the z-score quantifying the goodness-of-fit between the HRF (blue line) and the data (grey line).

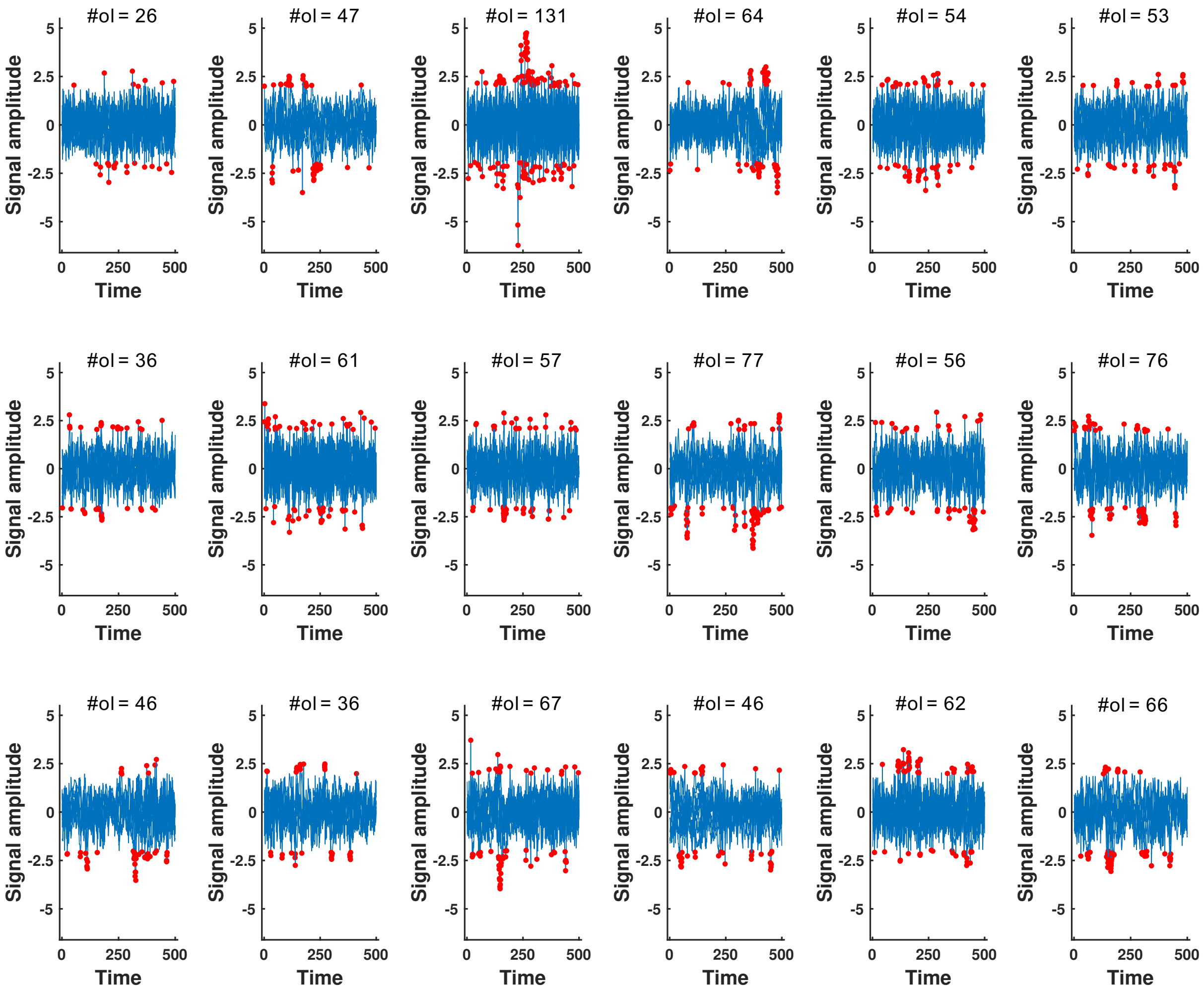


**Supplementary Figure 2: Resting-state network timeseries and outliers.** The plots are displayed in identical order to Figure 1 in the main text. In each plot, the timeseries of all nine resting-state networks are plotted superimposed (blue), with the x-axis representing time in volumes (500 in total). The y-axis represents the network timeseries signal amplitudes. Each individual network timeseries has been independently zero-centred around its median and scaled to have unit interquartile range, resulting in all outliers, highlighted in red, having values greater than ± 2. Above each plot is the total outlier count (#ol = number of outliers) across all networks.

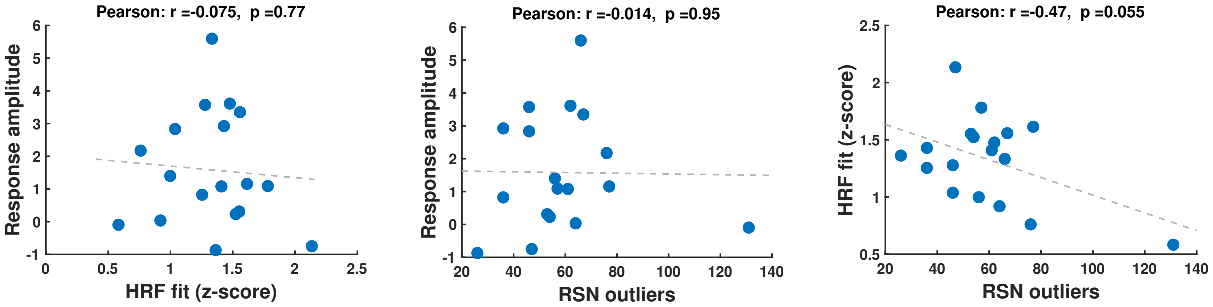


**Supplementary Figure 3: HRF fits, RSN outliers, and noxious-response amplitudes.** For all plots, the dashed grey line is the least squares line, and the Pearson correlation coefficient (r) and the associated p-value (p) are displayed overhead. Left: The correlation between noxious-response amplitude (y-axis) and noxious-response HRF goodness-of-fit (x-axis). Middle: The correlation between noxious-response amplitude (y-axis) and resting-state network timeseries outlier count (x-axis). Left: The correlation between noxious-response HRF goodness-of-fit (y-axis) and resting-state network timeseries outlier count (x-axis). Abbreviations: HRF = haemodynamic response function; RSN = resting-state network.

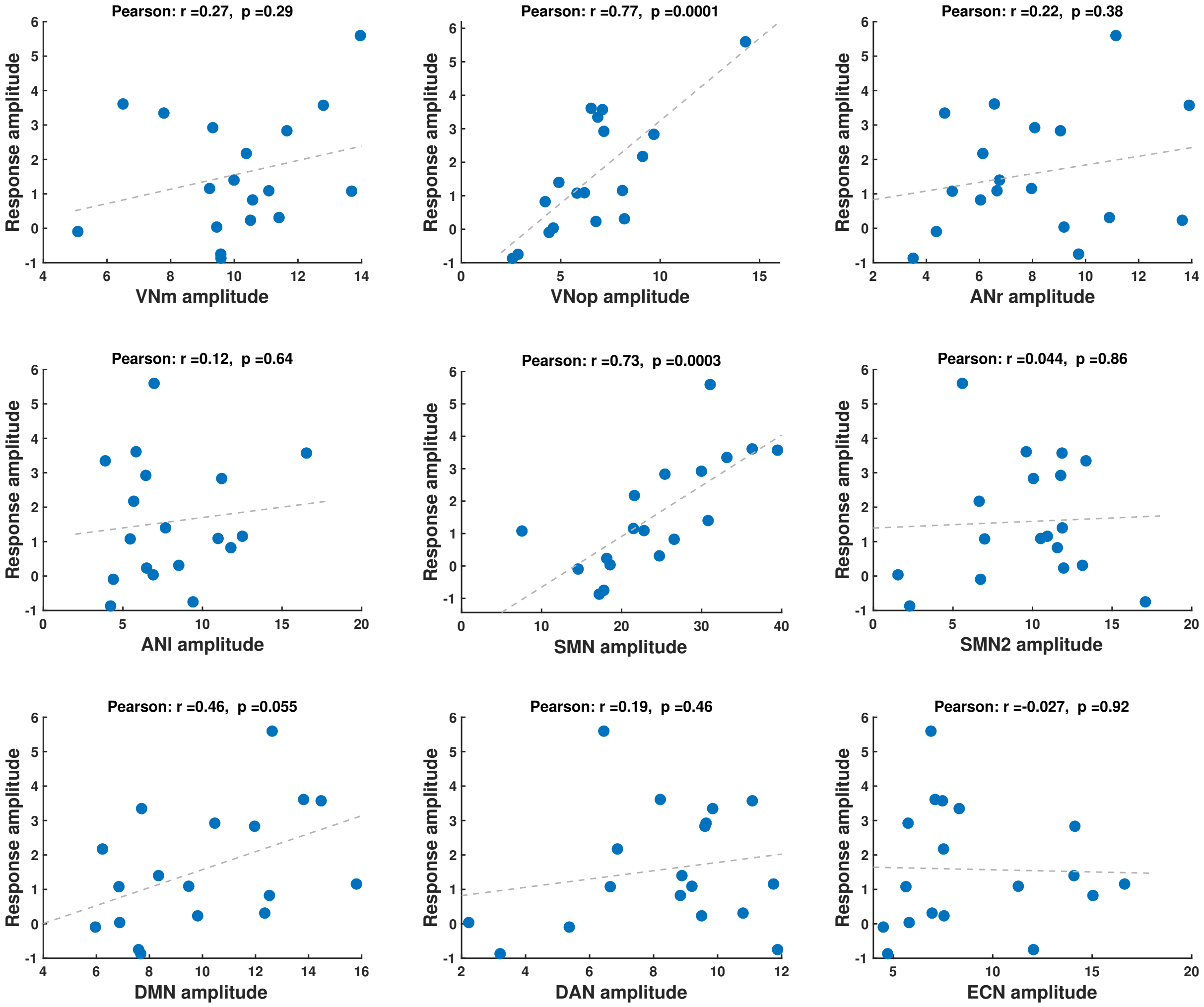


**Supplementary Figure 4: Noxious-response amplitudes vs resting-state network amplitudes.** The plots are displayed from left-to-right then top-to-bottom in identical order to that presented from left-to-right in Figure 2 in the main text. For all plots, the x-axis is the resting-state network amplitude, the y-axis is the noxious-response amplitude, the dashed grey line is the least squares line, and the Pearson correlation coefficient (r) and the associated p-value (p) are displayed overhead. Abbreviations: VNm = medial visual network; VNop = occipital pole visual network; ANr = right auditory network; ANl = left auditory network; SMN = somatomotor network; DMN = default mode network; DAN = dorsal attention network; ECN = executive control network.

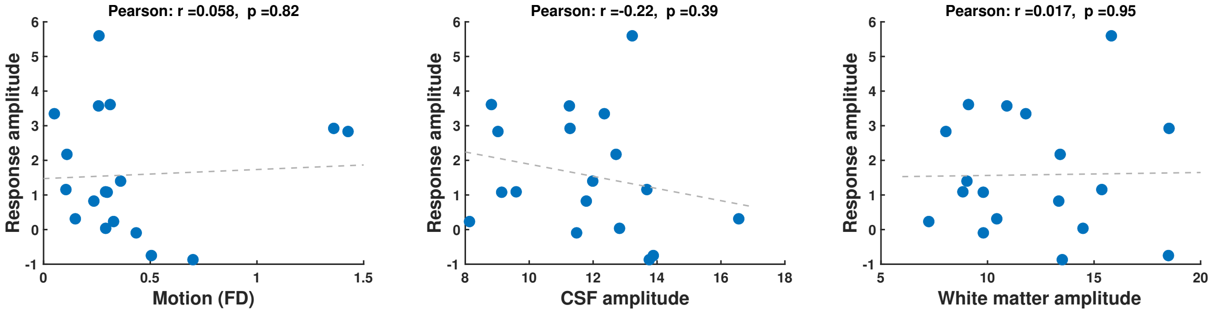


**Supplementary Figure 5: Noxious-response amplitudes vs resting-state imaging confounds.** For all plots, the x-axis is the resting-state imaging confound, the y-axis is the noxious-response amplitude, the dashed grey line is the least squares line, and the Pearson correlation coefficient (r) and the associated p-value (p) are displayed overhead. Abbreviations: FD = framewise displacement; CSF = cerebrospinal fluid.

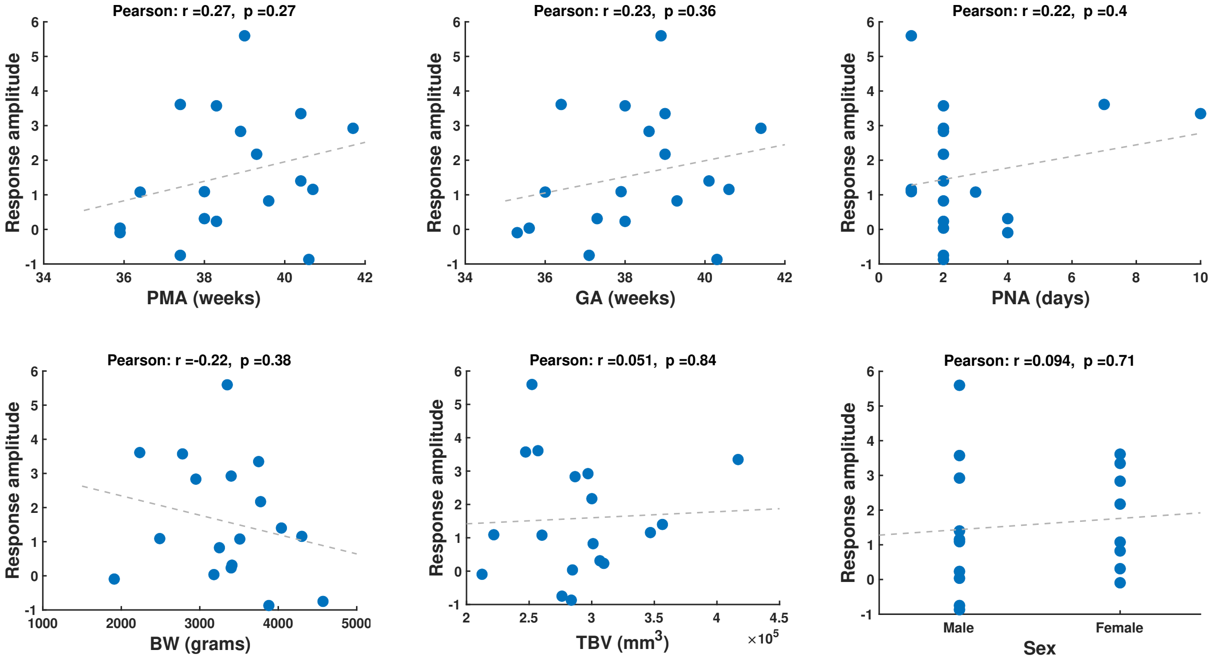


**Supplementary Figure 6: Noxious-response amplitudes vs clinical variables.** For all plots, the x-axis is the clinical variable (see Table 2 in main text), the y-axis is the noxious-response amplitude, the dashed grey line is the least squares line, and the Pearson correlation coefficient (r) and the associated p-value (p) are displayed overhead. Abbreviations: PMA = postmenstrual age; GA = gestational age; PNA = postnatal age; BW = birth weight; TBV = total brain volume

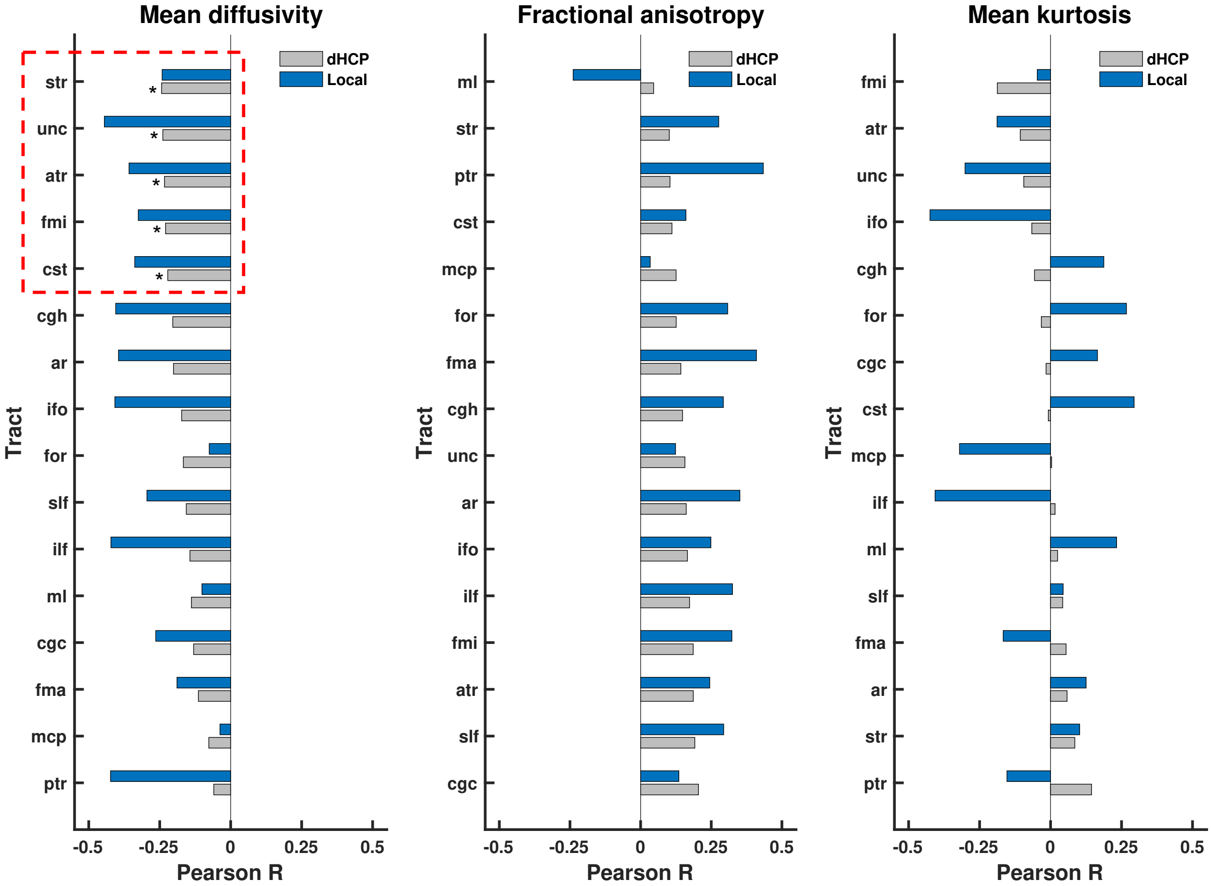


**Supplementary Figure 7: Correlations between noxious-response amplitudes and microstructural features of white matter tracts.** The three plots display the Pearson correlation coefficients (x-axis) between noxious-response amplitudes and microstructural features for each white matter tract (y-axis). For each tract, the coefficients for both the dHCP dataset (grey) an nociception-paradigm dataset (blue) are juxtaposed. The white matter tracts (y-axis) are independently ordered per plot according to the dHCP correlation coefficients: from most negative (top) to most positive (bottom). The asterisks and dashed red box indicates the five dHCP correlation coefficients identified as statistically significant. Abbreviations: ar = acoustic radiation; atr = anterior thalamic radiation; cgc = cingulate gyrus part of the cingulum; cgh = parahippocampal part of the cingulum; cst = corticospinal tract; fma = forceps major; fmi = forceps minor; for = fornix; ifo = inferior fronto-occipital fasciculus; ilf = inferior longitudinal fasciculus; mcp = middle cerebellar peduncle; ml = medial lemniscus; ptr = posterior thalamic radiation; slf = superior longitudinal fasciculus; str = superior thalamic radiation; unc = uncinate fasciculus.

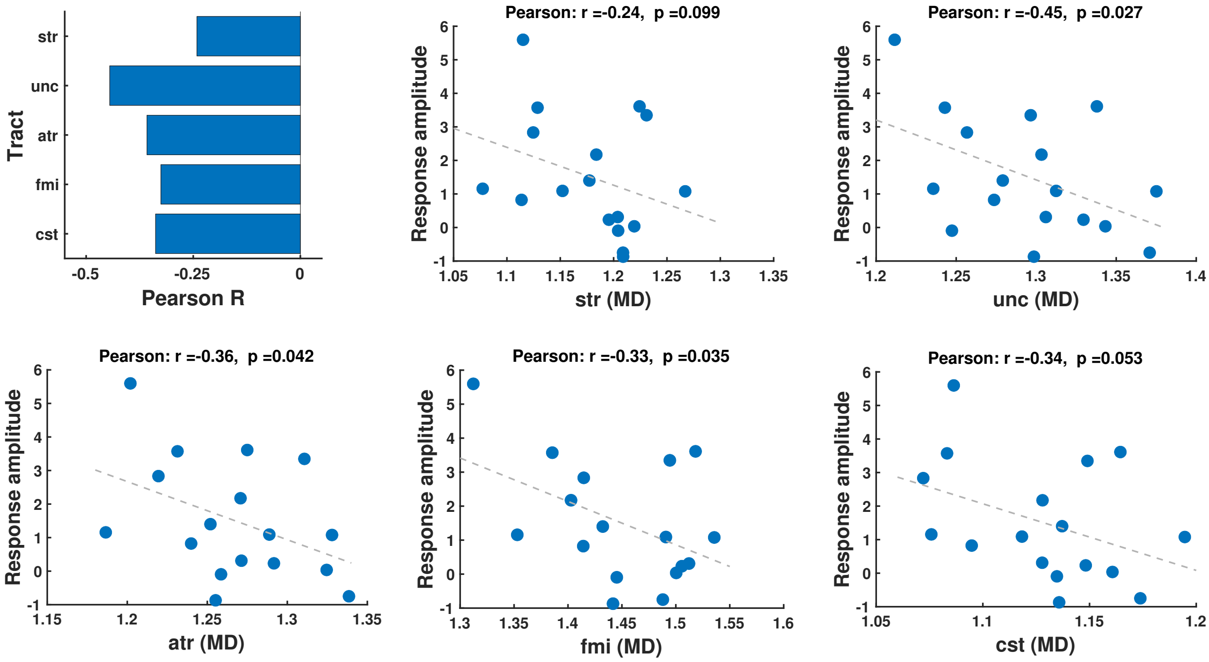


**Supplementary Figure 8: Nociception-paradigm dataset structure-function correlations for tracts identified in dHCP dataset.** All data presented are from the nociception-paradigm dataset only. The bar plot (top left) displays the Pearson correlation coefficients (x-axis) between noxious-response amplitude and mean diffusivity (MD) for the five tracts highlighted in the red box in Supplementary Figure 7. The scatter plots display the univariate correlations between noxious-response amplitudes (y-axis) and the MD of each white matter tract (x-axis) expressed in $\mu m^2$/ms. For all scatter plots, the dashed grey line is the least squares line, and the Pearson correlation coefficient (r) and the associated p-value (p) are displayed overhead. Abbreviations: atr = anterior thalamic radiation; cst = corticospinal tract; fmi = forceps minor; str = superior thalamic radiation; unc = uncinate fasciculus.

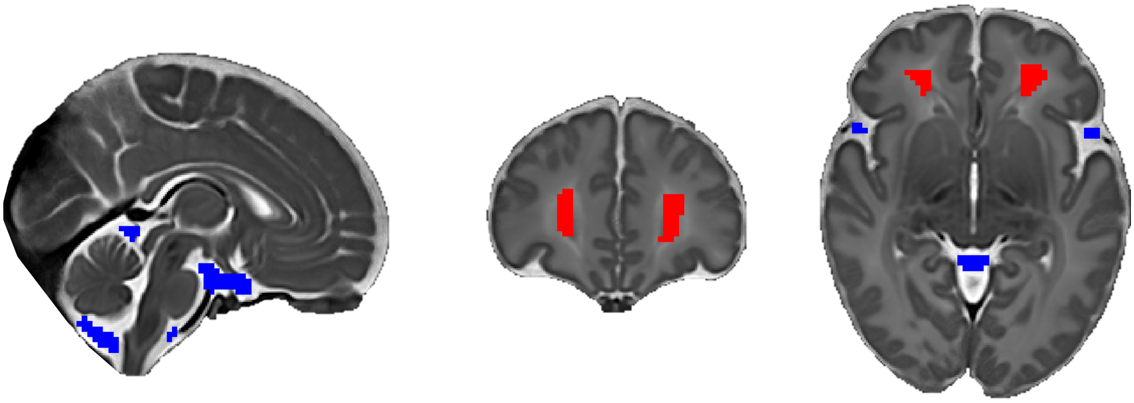

**Supplementary Figure 9: CSF and white matter ROIs.** The CSF ROI (blue) is restricted predominantly to the fluid surrounding the brainstem and cerebellum. The white matter ROI (red) is restricted to the largest white matter regions in the cerebrum tightly localized to the centre of white matter regions. Abbreviations: CSF = cerebrospinal fluid; ROI = region of interest.
